# Longitudinal multimodal characterization of radiation dermatitis in the C57BL/6J mouse model

**DOI:** 10.1101/2025.11.08.687376

**Authors:** Jasmine H. Wong, Sharen Rivas, Li Liu, Wade Koba, Patrik Brodin, Chandan Guha, Alana Deutsch, Adnan Mir, Beth N. McLellan, Kosaku Shinoda

## Abstract

Radiation dermatitis (RD) is a frequent and therapy-limiting toxicity of cancer radiotherapy, yet its mechanistic basis remains poorly understood. To address the need for quantitative, mechanistic insight, we developed a longitudinal C57BL/6J mouse model integrating clinical grading, non-invasive biophysical (“mesoscopic”) measurements, histopathology, and bulk transcriptomics. This controlled design enabled pseudo-longitudinal analysis of inflammation, barrier dysfunction, and fibrosis across four weeks after a single 25 Gy exposure. Transcriptomic profiling revealed early activation of phagosome formation, MHC I antigen presentation, and neutrophil degranulation preceding visible skin injury, followed by enrichment of IL-1, IL-10, and IL-13 signaling at peak disease. Mesoscopic erythema, fibrosis, and transepidermal water loss each displayed distinct temporal and molecular signatures, with limited overlap among gene sets. Histologic scoring confirmed early epidermal and inflammatory responses, linked to extracellular matrix remodeling and vascular development. The observed pathways align with prior studies implicating IL-17 and a senescence-associated IL-6/CCR6 axis, extending these findings into a quantitative, time-resolved framework. This multimodal resource bridges molecular, morphological, and functional scales, providing a platform for mechanistic and therapeutic studies of RD.

## Introduction

Radiation dermatitis (RD) is a common adverse reaction to radiation therapy (RT), affecting up to 95% of cancer patients.^1^ RD is categorized as acute, characterized by symptoms of erythema, desquamation, ulceration, and pigmentary changes, or chronic, demonstrating fibrosis and telangiectasia formation.^2^ These cutaneous symptoms can hinder adherence to the oncologic treatment plan and significantly affect quality of life. Despite its prevalence, RD’s pathophysiology is complex and remains poorly understood. Our current understanding of the mechanism involves the impairment of mitotic activity of progenitor cells from radiation-induced DNA damage, leading to a decrease in epidermal keratinocyte renewal and an increase in pro-inflammatory mediators, tissue damage, and oxidative stress.^3,4^

Common treatments include emollients, topical corticosteroids, or protective wound dressings,^5^ which are non-specific strategies that broadly reduce inflammation and protect the skin barrier. With an enhanced mechanistic understanding of this condition, it may be possible to utilize or design targeted treatments that optimize preventative and therapeutic management. Prior animal model studies of RD are scarce, variable in methodology, and limited to outcomes such as clinical grading and histology.^6,7^ To address this gap, we aimed to develop a temporal murine model of RD using clinical grading, objective assessments, histologic grading, and genome-wide gene expression analysis.

## Results

### Building a longitudinal multimodal dataset on radiation dermatitis (RD)

Because few animal studies have examined RD, its pathophysiology remains poorly understood. In humans, longitudinal tracking of RD onset and progression using tissue samples has clinical and ethical limitations. We established a murine multimodal dataset combining clinical grading, histologic analysis, objective skin parameter measurements (termed “mesoscopic” measurements in this study), and genome-wide gene expression. We used inbred C57BL/6J mice, which are genetically homogeneous, to effectively model repeated within-subject sampling in humans. Mice were divided into five groups—sham (no RT) and four post-RT groups—with weekly assessments over four weeks. At each time point, collected data included clinical photographs, mesoscopic measurements of erythema, fibrosis, transepidermal water loss (TEWL), and 4-mm punch biopsies from both irradiated skin and untreated skin on the contralateral flank (**Fig. 1A**). Radiation groups received a single 25 Gy dose to the left flank on day 0 using our novel irradiation setup (**Fig. 1B**). This dose was chosen based on prior studies demonstrating that 25 Gy reliably induces cutaneous changes resembling human RD.^8^

**Fig. 1.**
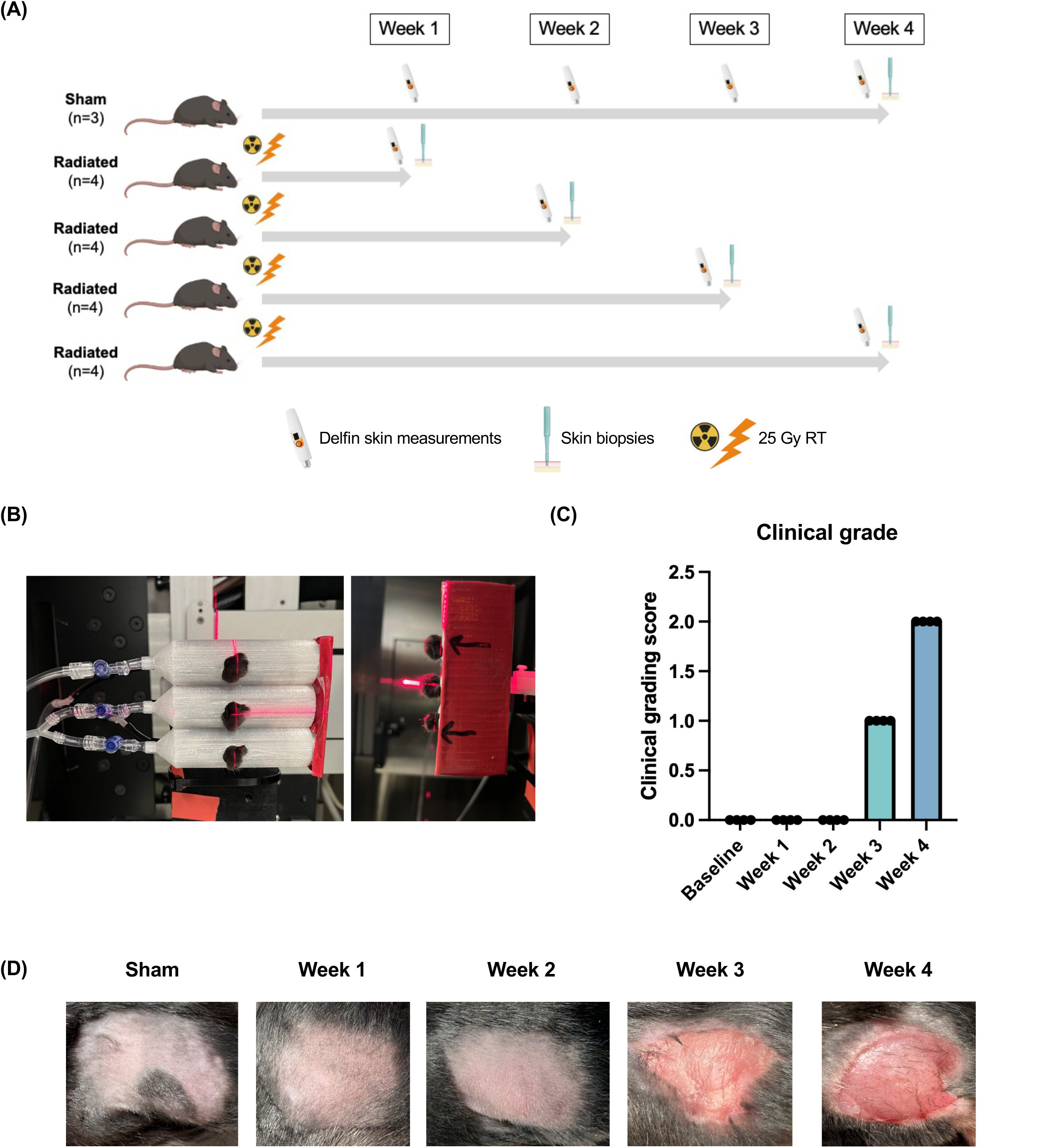
Longitudinal multimodal characterization of radiation dermatitis using a C57BL/6 model: **(A)** Schematic of experimental design with a single dose of 25 Gy radiation and data collected weekly for four weeks including mesoscopic measurements (Delfin) and skin punch biopsies. **(B)** Radiation setup with three mice at a time placed into 3D-printed cylindrical tubes with a circular cutout to isolate skin over the left flank for radiation and a 6mm thick lead shield on the rear of the jig to protect the animals from lethal exposure. **(C)** Clinical grading of radiation dermatitis severity, scored from 1-4, across four weeks. **(D)** Representative images of the progression of RD on the shaved left flank region across four weeks.

Clinical grading scoring was derived from human RD criteria, reflecting erythema and desquamation severity (scored 1-4 as described in the Methods), remained at zero through weeks 1-2, increased to 1 by week 3, and reached 2 by week 4, with consistent results across all mice (**Fig. 1C**). **Figure 1D** demonstrates photographic evidence of this progressive skin damage, closely resembling the morphology of human RD.^2^

### Early transcriptomic shifts precede clinical manifestation of RD

To identify the molecular mechanisms underlying the clinical phenotype, we performed RNA-sequencing on the skin biopsies and compared transcriptomic profiles of the irradiated and sham groups at each time point. Strikingly, based on a cut-off at log2fold>1 and p-value <0.05, more than 100 genes were differentially expressed at week 1 (72 downregulated, 37 upregulated; **Fig. 2A**). This substantial transcriptomic shift occurred before any clinical signs of RD. Gene Ontology (GO) analysis revealed early activation of phagosome formation, MHC I antigen presentation, and neutrophil degranulation, indicating that innate immunity was engaged well before overt clinical manifestations. By week 2, the number of DEGs increased to 253 (120 downregulated, 133 upregulated; **Fig. 2B**). Although the RD clinical grading score remained zero in the irradiated and sham groups, GO analysis revealed upregulation of Interleukin (IL)-10, IL-4, and IL-13 signaling, along with TREM1 and S100, proinflammatory mediators implicated in cutaneous inflammatory disorders such as atopic dermatitis and psoriasis (**Fig. 2B**). ^9–11^ By week 3, the number of DEGs peaked at 951 (309 downregulated, 642 upregulated; **Fig. 2C**), with strong enrichment of pathways related to neutrophil degranulation, pathogen-induced cytokine storm, and phagosome formation, a pronounced inflammatory surge. At week 4, the number of DEGs declined to 793 (237 downregulated, 556 upregulated; **Fig. 2D**) but remained enriched for similar inflammatory pathways, indicating persistent immune activation. Overall, transcriptomic profiling revealed a time-dependent expansion of DEGs from week 1 through week 3, followed by a modest decline at week 4 (**Fig. 2A–D**). This rise in DEGs indicates that radiation injury drives progressively broader transcriptional reprogramming, in parallel with the clinical progression of RD. These findings establish a foundation for identifying gene sets associated with clinical features of RD, which are examined in the following sections.

**Fig. 2.**
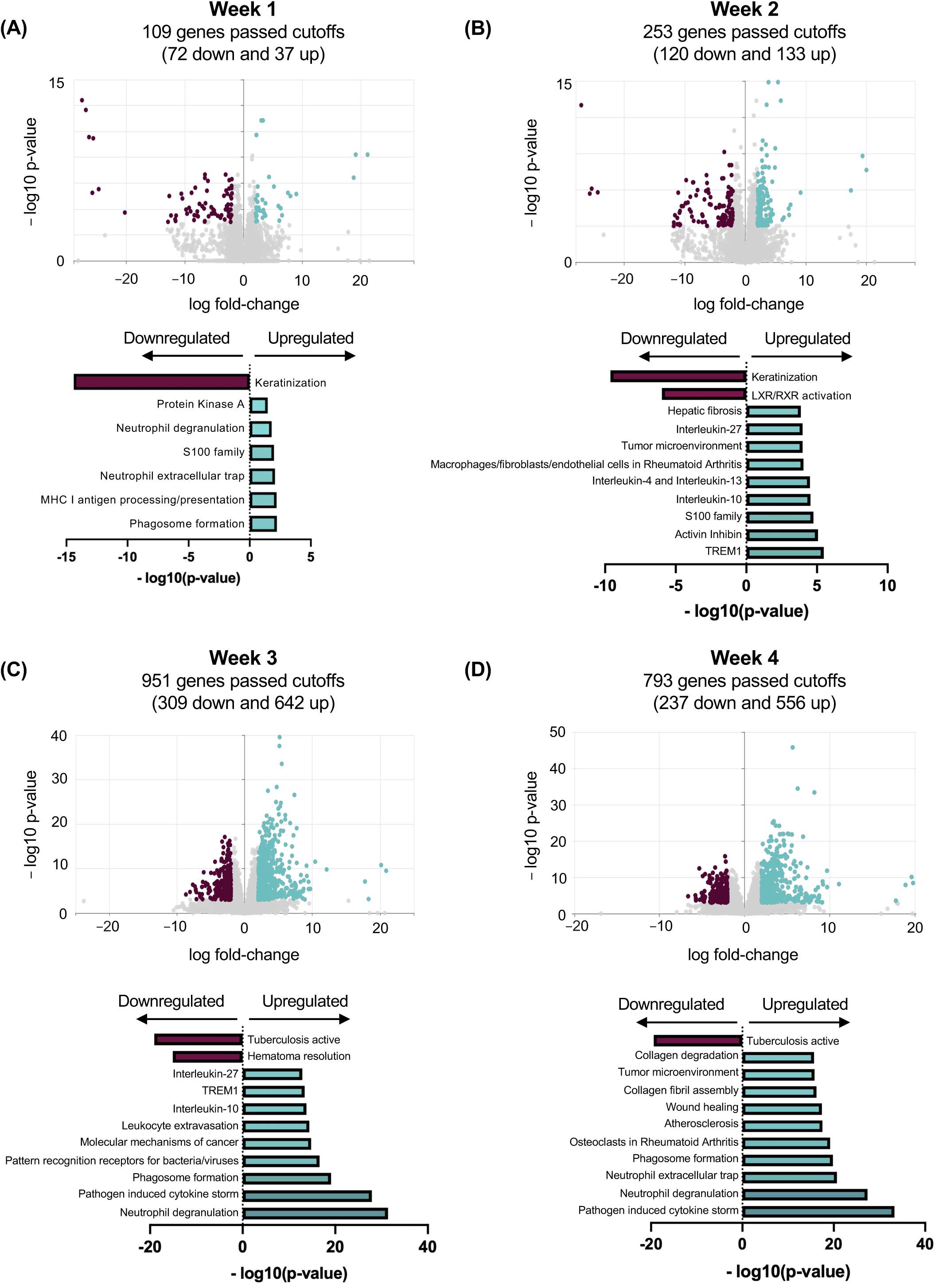
Time-dependent transcriptomic changes in the skin of mouse radiation dermatitis: **(A-D)** Volcano plots depicting the significantly downregulated and upregulated differentially expressed genes (DEG) and corresponding horizontal bar plots of Metascape pathway enrichment analysis depicting the most significantly downregulated and upregulated pathways associated with those DEGs at Week 1 **(A)**, Week 2 **(B)**, Week 3 **(C)**, and Week 4 **(D)**. Downregulated genes and pathways shown in red, upregulated genes and pathways shown in green, and genes that did not pass the cutoff of *** in grey.

### Mesoscopic parameters reveal three transcriptomic axes of RD

To bridge macroscopic (clinical) and microscopic (histologic) findings, we used Delfin Technologies devices to measure mesoscopic parameters—fibrosis, erythema, and TEWL—as objective readouts of skin response to irradiation. These parameters exhibited distinct temporal patterns (**Fig. 3A**). Erythema closely paralleled the clinical score, remaining unchanged relative to sham through week 2, then increasing gradually, and almost linearly, from week 3 to 4 (*P* < 0.001). Fibrosis, in contrast, showed no significant change through weeks 1–3 but rose abruptly at week 4, with the indentation force (in newtons) tripling compared to baseline (*P* < 0.0001). The week 4 values differed significantly from all earlier time points (*P* < 0.0001). Lastly, TEWL peaked at week 3. Levels were significantly higher at week 3 than baseline, week 1, and week 2 (*P* < 0.01), but declined by week 4 (*P* < 0.05). Notably, this decline coincided with the sharp rise in fibrosis, a reciprocal pattern which may reflect the transition from barrier disruption toward fibrotic remodeling. No significant differences from baseline were observed during weeks 1 and 2 for any of three parameters. Taken together, significant mesoscopic changes emerged later than transcriptomic alterations, aligning more closely with the onset of clinical symptoms around week 3. Importantly, each parameter displayed a unique temporal trajectory—gradual (erythema), abrupt (fibrosis), and transient (barrier dysfunction)—suggesting their distinct biological underpinnings and prompting further exploration of the gene signatures that correlate with each mesoscopic readout.

**Fig. 3.**
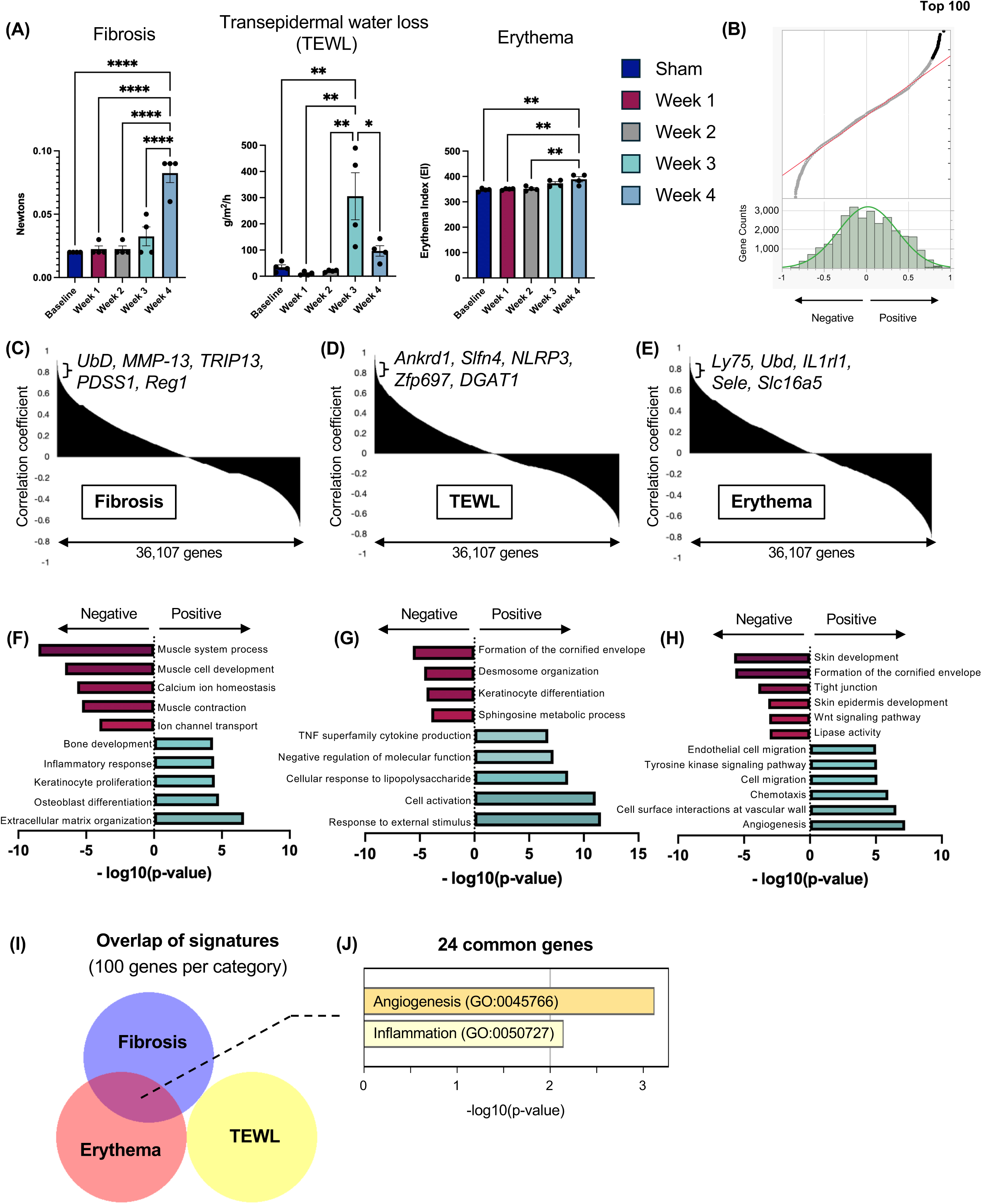
Mesoscopic assessment of radiation dermatitis progression and associated gene signatures: **(A)** Three mesoscopic variables (fibrosis, transepidermal water loss (TEWL), and erythema) scores across four weeks. Sham group in dark blue, Week 1 group in red, Week 2 group in grey, Week 3 group in teal, Week 4 group in blue. Statistical comparisons are based on Tukey’s multiple comparisons test performed in Prism (*p<0.05, **p<0.01, ****p<0.0001). **(B)** The Q-Q plot (quantile-quantile plot) comparing the top 100 genes with the null distribution. **(C-E)** Correlation coefficient analysis using Pearson correlations was conducted for all 36,107 genes and graphed for the three mesoscopic parameters – fibrosis **(C)**, TEWL **(D)**, and erythema **(E)** –highlighting the top five most significant genes. **(F-H)** Pearson correlations were ranked for all the genes and the top 100 genes with the highest correlations were determined, creating representative gene signatures for each variable. Horizontal bar plots of Metascape pathway enrichment analysis depicting the most significantly downregulated and upregulated pathways for fibrosis **(F)**, TEWL **(G)**, and erythema **(H)**. Downregulated pathways shown in red, upregulated pathways shown in green. **(I-J)** Overlap of the three mesoscopic gene signatures was assessed using a web application called BioVenn **(I-J)**. Fibrosis and erythema signature overlap demonstrated an overlap of 24 common genes, with pathways most significant for angiogenesis and inflammation.

To identify molecular correlates of mesoscopic skin changes, we ranked all genes by Pearson correlation with each parameter and selected the top 100 to identify representative gene signatures and putative molecular regulators. A threshold of 100 was chosen for consistency and interpretability, and Q-Q plots confirmed that these top genes deviated from the null, indicating non-random correlations (**Fig. 3B**). *Sele*, the gene encoding E-selectin (**Fig. 3E**) is one of the top molecular correlates of mesoscopic erythema. E-selectin is an endothelial adhesion molecule consistently reported to be upregulated in inflammatory dermatoses such as atopic dermatitis and psoriasis.^12,13^ Its correlation here highlights vascular activation as a shared mechanism across diverse skin inflammatory phenotypes. At the pathway level, erythema signatures were enriched for pathways related to angiogenesis, chemotaxis, and endothelial migration, reflecting vascular and inflammatory drivers of skin redness (**Fig. 3H**). For mesoscopic fibrosis, *UbD* and *Mmp13* were the two most positively correlated genes (**Fig. 3C**). *UbD* encodes ubiquitin D, which tags proteins for degradation and modulates signaling pathways such as mitogen-activated protein kinase (MAPK), Notch, and nuclear factor kappa-light-chain-enhancer of activated B cells (NF-κB), ultimately involved in regulating fibrogenesis.^14^ It also plays a critical role in collagen secretion by skin fibroblasts.^15^ Matrix metalloproteinase-13 (MMP13), a matrix metalloproteinase, is responsible for cleaving extracellular matrix (ECM) proteins, thereby facilitating fibrillar collagen remodeling. MMP3 is expressed by fibroblasts in human cutaneous wounds.^16^ Consistent with these functions, the most significantly enriched pathway in the fibrosis signature was ECM organization (**Fig. 3F**). For TEWL, *Nlrp3* and *Dgat1* were among the most strongly correlated genes (**Fig. 3D**). NLR family pyrin domain-containing 3 (NLRP3) promotes Interleukin-1β production, amplifying inflammatory responses, and has been found to be essential to the pathogenesis of rosacea.^17,18^ Diacylglycerol O-Acyltransferase 1 (DGAT1) catalyzes triacylglycerol synthesis and maintains retinoid homeostasis, both critical for preserving skin barrier integrity and preventing TEWL.^19,20^ Consistent with these roles, TEWL correlates showed downregulation of pathways related to cornified envelope formation, keratinocyte differentiation, and desmosome organization, reflecting barrier disruption (**Fig. 3G**).

Because fibrosis, erythema, and barrier disruption may arise from either shared or distinct mechanisms, we assessed the overlap among fibrosis, erythema and TEWL signatures. The three signatures were largely non-overlapping, underscoring that each mesoscopic parameter is dictated by a distinct molecular mechanism. However, fibrosis and erythema did share 24 (i.e., 24%) genes, enriched for angiogenesis and inflammatory pathways (*P* < 0.001), suggesting common vascular-inflammatory drivers. In contrast, the TEWL signature showed zero overlap with fibrosis or erythema (**Fig. 3I, J**). Together, these mesoscopic gene signatures not only delineate distinct and overlapping mechanisms of RD but as shown in the following section, provide a framework for high-resolution insights into biology of RD. The top 100 genes and their weekly fold changes are listed in **Suppl. Table 1.**

### Histologic scoring identifies early microscopic changes and their gene correlates

Skin sections from the same mouse cohort were evaluated by a dermatopathologist using a blinded scoring system (see Methods) to assess time-course histologic changes. Four parameters—epidermal change, fibrosis, inflammation, and follicular change—were graded on a 0–2 scale, capturing key microscopic hallmarks of RD. Representative histologic images illustrating the progression of radiation-induced changes across four weeks are shown in **Fig. 4A**, and representative images of the four variables are shown in **Fig. 4B**. Overall, all histologic variables followed trends similar to erythema, generally increasing in severity over the four weeks, except epidermal change, which rose at week 1, dipped at week 2, and then increased again through week 4. Notably, the acute response mirrored the transcriptomic finding, with robust increases in scores already evident at week 1 relative to the sham group, well before changes were detectable by clinical photographs or mesoscopic devices. This early rise was consistently observed across all four histologic parameters. Motivated by this similarity between transcriptomic and histologic dynamics, we performed a Pearson correlation analysis to identify putative factors driving these histologic changes.

**Fig. 4.**
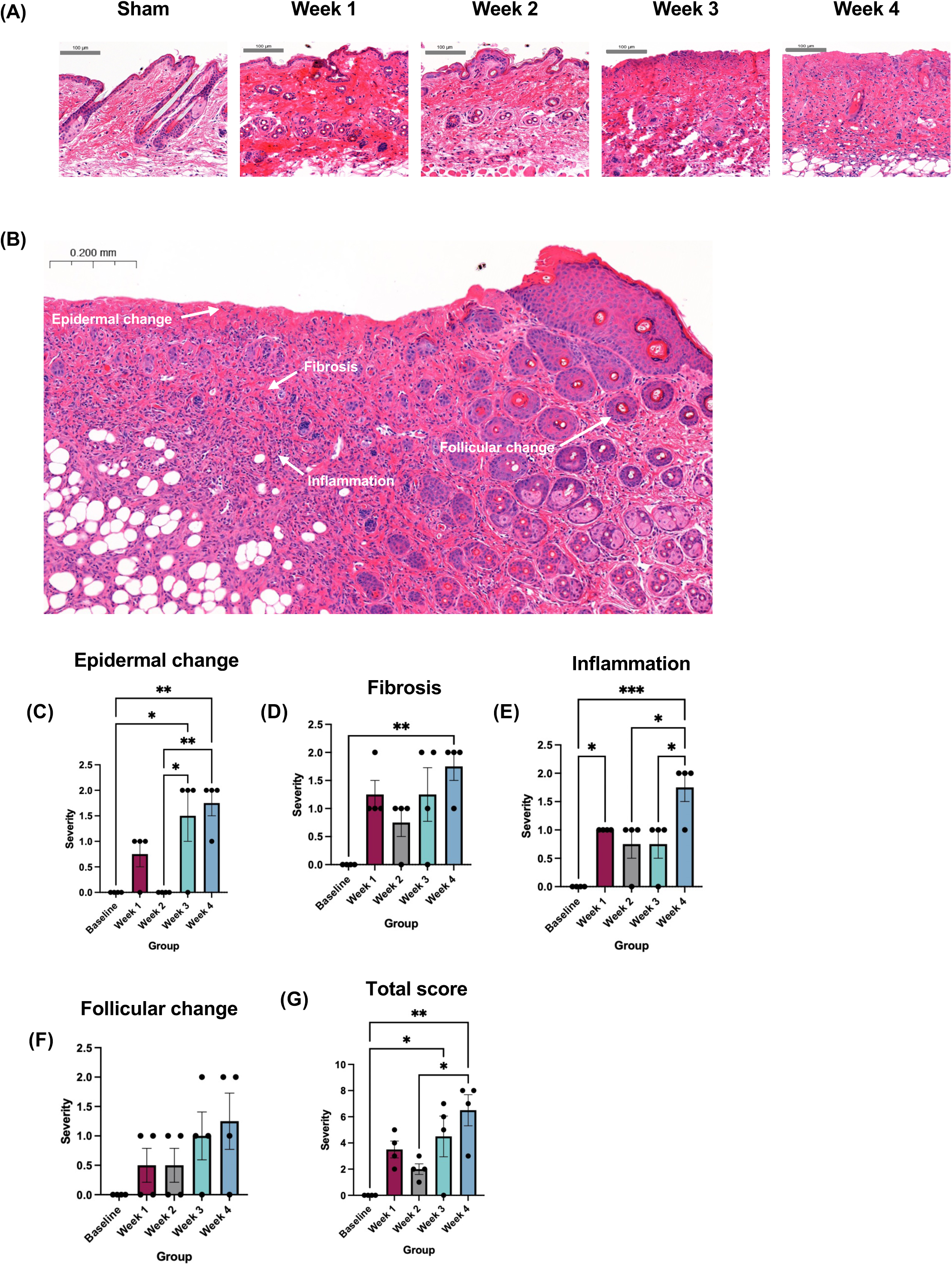
Histologic assessment of radiation dermatitis progression: **(A)** Representative hematoxylin and eosin (H&E)-stained histologic images of radiation-induced skin changes over the four weeks. **(B)** Representative H&E-stained histologic image depicting the four assessed variables (epidermal change, fibrosis, inflammation, and follicular change). **(C-G)** Four histologic variables for epidermal change **(C)**, fibrosis **(D)**, inflammation **(E)**, and follicular change **(F)** (graded on a 0-2 scale) and total score **(G)** (sum of four variables with a maximum score of 8) with scores graphed across four weeks. Blinded scores for histology variables with severity defined as: epidermal change (0=absent, 1=epidermal thickening or hyperkeratosis, 2=epidermal ulceration), fibrosis (0=absent, 1=moderate, 2=complete), follicular change (0=absent, 1=follicular dysplasia, 2=follicular loss), and inflammation (including dermal inflammation and hypodermal inflammation, 0=absent, 1=moderate, 2=complete). Sham group in dark blue, Week 1 group in red, Week 2 group in grey, Week 3 group in teal, Week 4 group in blue. Statistical comparisons are based on Tukey’s multiple comparisons test performed in Prism (\**P* < 0.05, ** *P* < 0.01, *** *P* < 0.001).

As with the mesoscopic parameters (**Fig. 3**), the top 100 genes for each variable were ranked by Pearson correlation to generate representative histological gene signatures. As was the case with the mesoscopic analyses, the top 100 correlates deviated to a significant extent from the null distribution via Q-Q analysis (not shown). Pathway enrichment was then assessed using Metascape. For epidermal change, none of the top five pathways were directly relevant (**Fig. 5A**), yet pathway analysis revealed downregulation of skin development, tight junction, and cornified envelope pathways, consistent with impaired barrier integrity. Notably, epidermal change was the only histologic variable lacking significant enrichment of pathways related to the ECM. This likely reflects tissue composition, as the epidermis is primarily composed of keratinocytes and relatively deficient in ECM compared to the ECM-rich dermis.^21^ For histologic fibrosis, *Adamts5* and *Abi3bp* emerged as the most positively correlated genes (**Fig. 5B**). The association of *Adamts5* with fibrosis is consistent with prior reports implicating this protease in dermal repair through fibroblast maturation,^22^ and it is also highly correlated with histologic inflammation in our dataset. *Abi3bp*, although less studied, has been suggested to promote mesenchymal stem cell proliferation, an essential process for cutaneous wound healing, highlighting it as a potentially novel contributor to cutaneous fibrosis. ^23^ As expected, fibrosis signatures were strongly enriched for ECM organization, collagen metabolic processes, and circulatory system pathways, consistent with fibrotic dermal remodeling, while showing downregulation of skin and gland development pathways (**Fig. 5G**). Notably, there was no overlap between the top 100 genes for mesoscopic fibrosis and histologic fibrosis, suggesting that these readouts capture distinct events. For histologic inflammation, *Plat* and *Kng2* were the most highly correlated genes (**Fig. 5C**). *Plat*, a plasminogen activator, has long been implicated in inflammatory skin disorders such as psoriasis, and plasminogen-deficient mice display impaired wound healing.^24,25^ *Kng2*, which participates in kinin formation and keratinocyte differentiation, is proinflammatory; kinins have been shown as elevated in dermatoses such as psoriasis and atopic dermatitis.^26^ Together, these associations highlight the importance of plasminogen and kinin pathways in cutaneous inflammation. Inflammation signatures showed enrichment for complement and coagulation cascades, ECM organization and proteoglycans, and vascular development (**Fig. 5H**). Finally, for follicular change, we identified *Arpc5*, a relatively uncharacterized actin filament–related gene (**Fig. 5D**).^27^ Although not directly linked to follicular biology, knockout of the actin related protein 2/3 (Arp2/3) complex in the murine epidermis produces severe psoriasis-like phenotypes, suggesting a potential role in skin homeostasis. Furthermore, follicular change was associated with pathways related to metabolic processes, including hydroxy and indole compound metabolism, as well as membrane organization (**Fig. 5I**). The total histologic score integrated these findings, demonstrating pathway enrichment for ECM organization, vasculature development, and gland development, capturing the combined fibrotic, vascular, and epidermal contributions to RD (**Fig. 5J**). The top 100 genes and their weekly fold changes are listed in **Suppl. Table 2.**

**Fig. 5.**
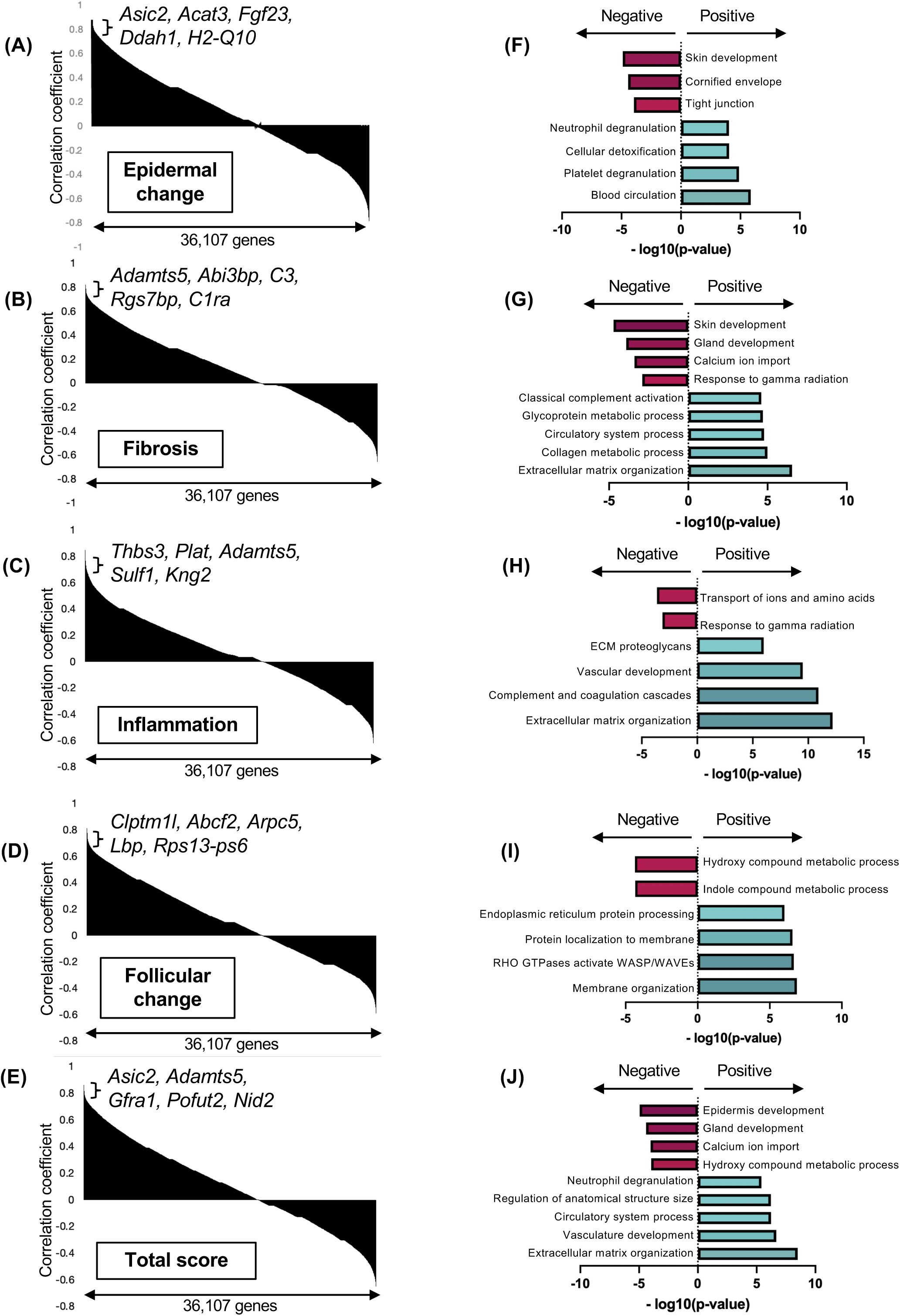
Identification of gene signatures and pathways associated with each histologic feature: **(A-E)** Correlation coefficient analysis using Pearson correlations was conducted for all 36,107 genes and graphed for the five histologic parameters –epidermal change **(A)**, fibrosis **(B)**, inflammation **(C)**, follicular change **(D)**, and total score **(E)** –highlighting the top five most significant genes. **(F-J)** Pearson correlations were ranked for all the genes and the top 100 genes with the highest correlations were determined, creating representative gene signatures for each variable. Horizontal bar plots of Metascape pathway enrichment analysis depicting the most significantly downregulated and upregulated pathways for epidermal change **(F)**, fibrosis **(G)**, inflammation **(H)**, follicular change **(I)**, and total score **(J)**. Downregulated pathways shown in red, upregulated pathways shown in green.

In comparing these histologic gene signatures against one another, the five signatures were largely unique and did not overlap with each other, except for the total score, as overlap is expected due to its composite nature (**Fig. 6**). The greatest overlap was observed between epidermal change and follicular change, with 11 shared genes (2.2% of the total gene set). A smaller overlap was noted between follicular change and fibrosis (seven genes, 0.3%). In contrast, inflammation showed minimal overlap with the other categories (20.2%), indicating a largely distinct gene signature. Taken together, these histologic analyses reveal that RD involves multiple, largely independent molecular programs corresponding to distinct microscopic hallmarks. Their partial convergence only at the composite score underscores the multifactorial and temporally heterogeneous nature of radiation-induced skin injury.

**Fig. 6.**
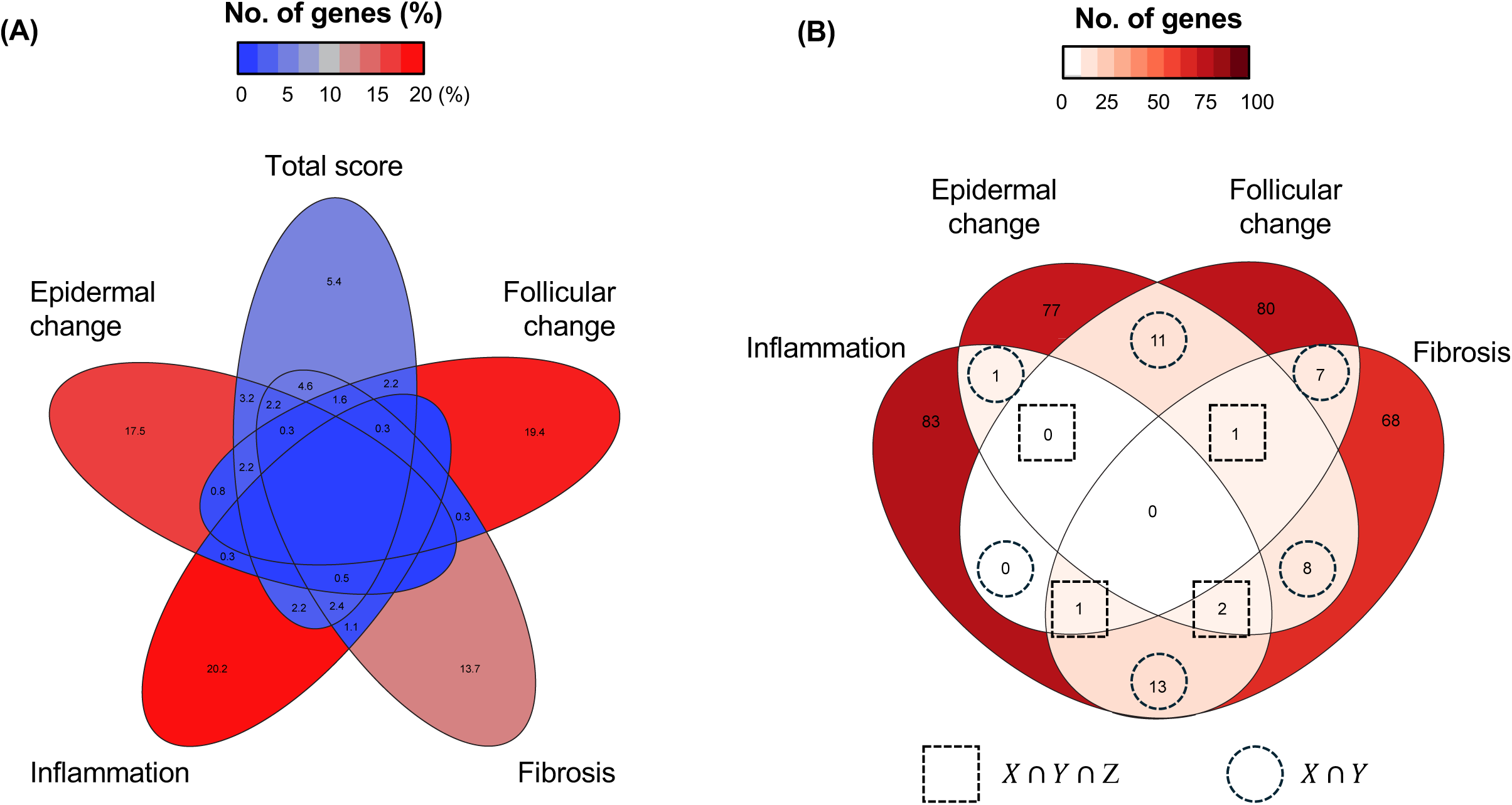
**(A)** Five-way Venn diagram showing percentage overlap between mouse histologic gene signatures and human radiation dermatitis gene signatures for epidermal change, follicular change, fibrosis, inflammation, and total score. Percentage values indicate proportion of mouse genes overlapping with each category. Color intensity (blue to red) reflects increasing percentage of overlapping genes. **(B)** Four-way Venn diagram displaying absolute number of overlapping genes across four gene signatures (epidermal change, follicular change, fibrosis, inflammation). Shaded color scale (light to dark red) corresponds to increasing number of genes. Dashed boxes denote triple overlaps, and dashed circles denote pairwise overlaps.

## Discussion

RD remains one of the most prevalent and impactful toxicities of cancer RT, yet molecular mechanistic insights have been limited. In this study, we established a multiscale, longitudinal dataset that systematically connects clinical, mesoscopic, histologic, and transcriptomic features of RD. To our knowledge, this is the first preclinical resource to couple quantitative, non-invasive biophysical measurements with matched histologic and genome-wide gene expression profiling that capture the temporal evolution of RD.

Our transcriptomic findings align with prior mechanistic studies elucidating immune drivers of RD. *Liao et al.* demonstrated that IL-17-expressing γδ T cells are essential mediators of radiation-induced skin inflammation in C57BL/6 mice, acting through IL-1β and IL-23 signaling and driving keratinocyte proliferation and acanthosis.^8^ The enriched pathways in our dataset—particularly neutrophil degranulation, phagosome formation, and IL-1/IL-13/IL-10 signaling—resonate strongly with these immune mechanisms, suggesting that early innate activation precedes and possibly amplifies the cytokine cascades previously observed in γδ T-cell-dependent models. Consistent with Liao’s findings, our GO analysis also highlights keratinocyte differentiation and desmosome organization as downregulated processes in irradiated skin, reinforcing the concept that epidermal barrier disruption and compensatory hyperproliferation are central to RD pathophysiology. Moreover, *Paldor et al.* extended this paradigm using single-cell transcriptomics to identify a senescence-associated IL-6/CCR6 axis that orchestrates IL-17-producing T-cell recruitment.^28^ When viewed together, these studies delineate a continuum of inflammatory signaling—from IL-1β and IL-23–driven γδ T-cell activation ^8^ to IL-6/CCR6-mediated immune recruitment ^28^—that underpins radiation-induced skin injury. Our bulk RNA-seq and temporal analyses integrate these mechanistic insights within a longitudinal, multiscale framework, revealing how innate immune activation, epidermal stress responses, and fibrotic remodeling unfold sequentially over time.

Our model leverages the genetic homogeneity of the C57BL/6J inbred strain, which minimizes inter-animal variability and effectively approximates repeated within-subject sampling—a condition challenging to achieve in humans. This genetically controlled framework enabled consistent temporal trajectories of disease progression across animals, revealing reproducible molecular correlates of erythema, fibrosis, and barrier disruption. Recently, *Lawrence et al.* employed a hairless SKH-1 mouse model that more closely mimics human skin architecture and pigmentation.^29^ Because SKH-1 is an inbred strain, applying a similar multimodal, genome-wide approach at larger scale in their system could yield complementary insights and deepen the human relevance of preclinical RD research. One limitation of our model is our use of a single dose of radiation rather than repeated fractionated doses. Future models should consider alternative RT dosing schedules as are often used in treatment of cancers.

In conclusion, this study establishes a reproducible and cost-effective multimodal platform for studying RD in C57BL/6J mice. By capturing the temporal interplay between inflammation, barrier dysfunction, and fibrosis across quantitative scales, it not only refines the understanding of RD pathogenesis but also provides a valuable open resource for the dermatologic and radiation biology communities.

## Materials and Methods

### Mouse irradiation setup

All procedures were performed in accordance with protocols approved by the Institutional Animal Care and Use Committee of the Albert Einstein College of Medicine. Ten-week-old female C57BL/6J mice (Jackson Laboratory, #000664) were randomly divided into sham (no RT) and irradiated groups. For radiation, three mice at a time were anesthetized with inhaled isoflurane and placed in custom 3D-printed cylindrical jigs that maintained continuous anesthesia and oxygen flow. A circular opening exposed the left flank for irradiation, while a 6-mm lead shield protected the remainder of the body. Irradiation was performed using an X-ray source (Xstrahl SARRP) operated at 220 kV peak voltage and 13 mA current in broad-beam mode, delivering a single 25 Gy dose over 420 seconds (3.6 Gy/min) at a 35 cm focus-to-skin distance (FSD). Details pertaining to the irradiation settings are described in the **Suppl. Note.** After treatment, the exposed area was shaved to visualize skin changes and marked for longitudinal assessment.

### Clinical grading

To score the severity of skin injury in the mouse model, a modified scoring system was developed based on the National Cancer Institute Common Terminology Criteria for Adverse Events (CTCAE version 5.0).^30,31^ Blinded clinical images were graded by a board-certified dermatologist from 1-4, with grade one indicating dry desquamation and minimal erythema, grade two indicating brisk erythema with patchy moist desquamation, grade three indicating severe moist desquamation, and grade four indicating ulceration, bleeding, or skin necrosis. Although none of our mice reached a severity threshold of greater than 2 with signs such as ulceration or skin necrosis, we chose this scoring system since it is more directly translatable to clinical grading for RD in humans. Additionally, future studies that utilize longer time courses or increased radiation doses that result in more severe RD may choose to utilize the same scoring system.

### Tissue histology

Skin tissue was fixed in 4% formaldehyde for 24 hours and transferred to 70% ethanol, then embedded in paraffin, sectioned with a vertical cross-section, and stained with hematoxylin and eosin (H&E) using a standard protocol. Blinded histologic images were graded by a dermatopathologist for following four variables on a 0-2 scale: epidermal change (0=absent, 1=epidermal thickening or hyperkeratosis, 2=epidermal ulceration), fibrosis (0=absent, 1=moderate, 2=complete), follicular change (0=absent, 1=follicular dysplasia, 2=follicular loss), and inflammation (including dermal inflammation and hypodermal inflammation, 0=absent, 1=moderate, 2=complete). These histologic categories were combined into a total score by calculating the total sum of the four categories, with a maximum score of eight.

### Non-invasive skin measurements

Objective measurements of erythema, fibrosis, and transepidermal water loss (TEWL) were deemed “mesoscopic” assessments, as in between our macroscopic (clinical grading) and microscopic (histologic) evaluations. For these mesoscopic variables, measurements were taken in triplicate and averaged. Erythema (SkinColorCatch), fibrosis (SkinFibroMeter), and TEWL (VapoMeter) were all measured using devices from Delfin Technologies. The SkinColorCatch quantifies relative red, green, and blue pigment in the skin, allowing differentiation between erythema and melanin. The SkinFibroMeter compares the force in Newtons required to compress an indenter against skin, while the VapoMeter assesses evaporation rate in grams/meters²/hour, a surrogate for skin barrier function.

### Statistics

Statistically significant differences between the mesoscopic assessments and histologic variable groups were assessed using one-way analysis of variance (ANOVA) with Tukey’s multiple comparisons test using Prism v10 (GraphPad) software. A p-value of less than 0.05 was considered significant.

### Bulk RNA-Sequencing

Four-millimeter punch biopsies from both the irradiated skin and the contralateral flank were collected and cut in half and used for histology and RNA-seq. Skin tissue was stored in RNAlater and frozen at -80 degrees Celsius until further processing. Total RNA was extracted using the RNeasy mini-kit (Qiagen). RNA samples were quantified using Qubit 2.0 Fluorometer (ThermoFisher Scientific, Waltham, MA, USA) and RNA integrity was checked with 4200 TapeStation (Agilent Technologies, Palo Alto, CA, USA). rRNA depletion sequencing library was prepared by using QIAGEN FastSelect rRNA HMR Kit (Qiagen, Hilden, Germany). RNA sequencing library preparation used NEBNext Ultra II RNA Library Preparation Kit for Illumina by following the manufacturer’s recommendations (NEB, Ipswich, MA, USA). Briefly, enriched RNAs are fragmented for 15 minutes at 94°C. First strand and second strand cDNA are subsequently synthesized. cDNA fragments are end repaired and adenylated at 3’ends, and universal adapters are ligated to cDNA fragments, followed by index addition and library enrichment with limited cycle PCR. Sequencing libraries were validated using the Agilent Tapestation 4200 (Agilent Technologies, Palo Alto, CA, USA), and quantified using Qubit 2.0 Fluorometer (ThermoFisher Scientific, Waltham, MA, USA) as well as by quantitative PCR (KAPA Biosystems, Wilmington, MA, USA). ERCC RNA Spike-In Mix (Cat: #4456740) was added to normalized total RNA prior to library preparation following manufacturer’s protocol. The sequencing libraries were multiplexed and clustered onto a flowcell on the Illumina NovaSeq instrument according to manufacturer’s instructions. The samples were sequenced using a 2×150bp Paired End (PE) configuration, targeting ∼30 million reads per sample. Image analysis and base calling were conducted by the NovaSeq Control Software (NCS). Raw sequence data (.bcl files) generated from Illumina NovaSeq was converted into fastq files and de-multiplexed using Illumina bcl2fastq 2.20 software. One mismatch was allowed for index sequence identification Raw fastq files were deposited in the Gene Expression Omnibus database and are available under accession #GSE300340.

### Bioinformatics

Bioinformatic analysis was performed using Kallisto and DESeq2 software. Bioinformatic analysis using Kallisto generated gene expression data across all timepoints for 36,107 genes. Fastq files were trimmed for adaptor and low-quality sequence using Trim Galore v0.6.7. Transcript-level counts were estimated by Kallisto v0.46.2 using mm10 reference genome. Outputs from Kallisto were imported into R using the Bioconductor package tximport v1.28.0, and the expression levels of each gene were estimated. Differential expression analysis was then performed using DESeq2 v1.40.2. Metascape, an online gene annotation and analysis resource, was used for pathway and process enrichment analysis of these 100 genes.^32^ A comparative analysis between human and mouse was conducted using human differentially expressed genes (DEG) and the mouse top 100 genes. For the cut-off to determine DEGs, log2fold≥1 and raw p-value <0.05 were used. QIAGEN Ingenuity Pathway Analysis (IPA) was used to identify canonical pathways and upstream regulators.^33^

### Multivariate statistics

Pearson correlation coefficient analysis was conducted in Microsoft Excel for the mesoscopic and histologic parameters measured. Correlations were ranked across 36,107 genes, and the top 100 with the highest coefficients were selected to generate representative gene signatures for each parameter. To assess whether these observed correlations reflected genuine associations with our parameters of interest rather than spurious correlations, quantile–quantile (Q–Q) plots were generated in JMP (SAS Institute). Gene signature overlaps were assessed using the BioVenn web application.^34^

### Power analysis

Following the framework of Wu et al.^35^ and Li et al.,^36^ simulations were conducted in R to estimate power across various effect sizes by resampling empirical distributions of gene expression levels and dispersions. For each gene, power was calculated individually rather than assuming a single common variance, as it varied widely across the transcriptome. Differential expression detection was modeled with DESeq2, and power was assessed at conventional thresholds (fold change ≥1.5 or ≥2.0; *P* < 0.05).

### Declaration of Interests

The authors have no interests to declare.

## Acknowledgements

This study was supported by NIH grants DK110426 (KS). The McLellan lab is supported by the Montefiore Einstein Comprehensive Cancer Center (MECCC) Interventional Investigator-Initiated Trials grant, the American Academy of Dermatology Hair Loss and Alopecia Initiative in Research (HAIR) grant, and the LEO Foundation (Copenhagen, Denmark). S.R. was supported by the American Academy of Dermatology–Society for Investigative Dermatology (AAD–SID) Gap Year Fellowship. The Shinoda lab gratefully acknowledges support from the Hevolution Foundation (*via* American Federation for Aging Research).

## Author contributions

J.H.W. and S.R. jointly performed mouse handling and sample collection. S.R. performed biopsy sample processing for RNA extraction. L.L. performed bioinformatic analyses of the RNA-seq dataset. W.K. designed and 3D-printed the radiation setup and administered the radiation dose to the mice. P.B. and C.G. designed the radiation setup and calculated the radiation dose. A.M. designed the histology variable grading system and conducted all histopathologic analysis. A.D. and B.N.M. designed the clinical and mesoscopic variable grading systems, conducted all clinical grading, directed the clinical study, and edited the manuscript. J.H.W. and K.S. conceived the project, created the figures, and wrote the paper with input from all the authors.

## Notes

https://www.ncbi.nlm.nih.gov/geo/query/acc.cgi?acc=GSE300340

### Competing Interest Statement

The authors have declared no competing interest.

