## Supplementary material for "Longitudinal multimodal characterization of radiation dermatitis in the C57BL/6J mouse model": Suppl. Note

**Supplementary Note**

**Radiation Parameters:**

X-ray Irradiator: Xstrahl's SARRP (Small Animal Radiation Research Platform)

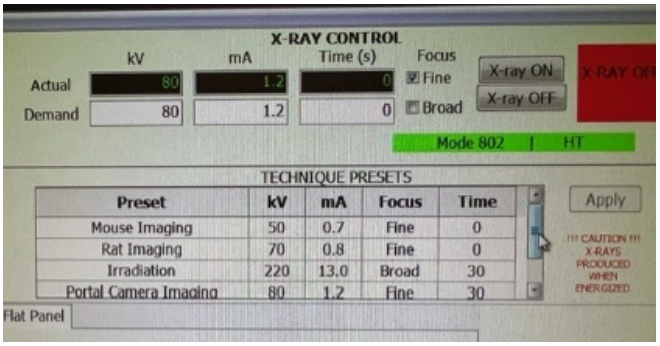


**SARRP Treatment Parameters:**

Maximum Current: 13mA

Maximum Voltage: 220kV at peak

Focus: Broad beam

Cu Filter dimensions: 0.15 mm

System output: 5Gy/minute

Focal length of beam: 35mm (Focus to Skin Distance)

Treatment time: 420 seconds

Total: 25 Gy

**SARRP QA and Jig setup on the SARRP:** Before starting any experiment, the SARRP robotics are checked to ensure all motion on X, Y, Z and couch are functional before proceeding with the warmup. After successfully homing the SARRP, either short or long warmup is selected for the tube conditioning. The "High Tension" is off, and the key is always on "Standby" when the interlocks are open and on "High Tension" when x-rays are being energized. To get the jig in place on the SARRP table, the couch is rotated 180 degrees then allowed to fully stop, proceeded by rotating the gantry arm to 90 degrees. The collimator is completely removed and a 0.15mm Copper filter is placed in the filter slot. The jig is specially designed to attached directly to the SARRP table and held on by a set screw to secure animals. Existing anesthesia lines and scavenger filter lines are connected to the jig to ensure the animals are continuously sedated and excess anesthesia is being collected to a charcoal filter.

**Animal Setup:** Individual animals were treated on a specialized three animal jig complete with anesthesia delivery during the injury. Mice were placed in the prone position and run off gas was scavenged by use of a charcoal filter. Skin with fur is gently pulled through the provided holes during the treatment. Once irradiated, animals are trimmed at the location at the hole by trimmer. This was continued up to three weeks until the hair started to fall off and physically demarcate the area.

Anesthesia was delivered to animals at 1.5% and 1.5ml/min, with animals subsequently induced in induction chamber for five minutes before placing mouse in the jig. The mice were marked with a permanent marker on the tail for tagging of completion. This was to ensure the animal was only treated once and not skipped. The existing anesthesia line of the system was used to keep the animals under sedation during the treatment part of the experiment.

**At 90 degrees:** The gantry was targeted, centered with lasers on the pulled skin/fur through the hole in the jig on all three mice before treatment commenced. There was a 6mm thick shield on the rear of the jig to protect the animals from lethal exposure.
